# HP6/Umbrea, a rapidly evolving *Drosophila* HP1-family paralog, is a candidate HP1a-recruited plasticizer of heterochromatin

**DOI:** 10.64898/2026.08.24.746899

**Authors:** UnJin Lee, Li Zhao

## Abstract

HP6/Umbrea, a rapidly evolving *Drosophila* (Heterochromatin Protein 1) HP1-family paralog, has well-documented sequence and regulatory evolution but an under-studied molecular function. In this manuscript, we hypothesize that HP6/Umbrea acts as an HP1-recruited plasticizer, providing support for this model using coarse-grained molecular-dynamics simulations of HP1a condensates. Retaining only the dimerizing chromoshadow domain (CSD), HP6/Umbrea notably lacks independent chromatin-binding capacity but binds HP1a directly, co-localizing with it *in vivo*. We report that when covalently tethered to an HP1a carrier, HP6/Umbrea partitions into HP1a condensates ∼6-fold more strongly than when free, supporting HP1a-mediated recruitment as its entry route. Once incorporated, HP6/Umbrea leaves the phase-separation threshold, interfacial tension, and host partitioning statistically unchanged, but monotonically lowers dense-phase density. These observations are consistent with a spacer function rather than generic loss of cohesion. Importantly, unchanged short-time internal mobility suggests a packing effect, predicting increased permeability to large transcriptional machinery, potentially resulting in a position effect-variegation (PEV)-like modulation of heterochromatic silencing. Finally, comparative sequence analysis shows the C-terminal tail is a recently originated, purifying-selection-constrained innovation, which is consistent with an evolved function in this region. In sum, our simulations suggest a mechanistic basis for how HP6/Umbrea may have evolved as a condensate plasticizer and thus potentially act as a rheostat for leaky transcription.

**Author summary:** Heterochromatin, the densely packed, gene-silencing fraction of the genome, is organized in part by Heterochromatin Protein 1 (HP1), which has been described as forming liquid-like condensates. *Drosophila* carries a fast-evolving duplicate of HP1, called HP6/Umbrea, which has been extensively studied as a young gene under strong selection. It is known to interact with HP1 and other heterochromatin proteins, yet a mechanistic account of what it does, especially the biochemical function that natural selection could act on, has remained lacking. HP6/Umbrea is a truncated protein that retains only the domain that lets HP1 proteins pair up, having lost the parts that read and bind chromatin. Using physics-based simulations of the HP1 condensate, our simulations indicate that this reduced structure has a simple consequence: HP6/Umbrea cannot enter heterochromatin on its own but is carried in by pairing with HP1, and once inside it loosens the interior packing without changing the condensate’s boundary or its tendency to form. We propose that HP6/Umbrea acts like a plasticizer, a molecular softener that could make otherwise silenced heterochromatic genes leakier (e.g., more widespread transcription) and thereby tune repression.

## Introduction

HP6, also called Umbrea, is a well-characterized young gene in *Drosophila*. A rapidly evolving Heterochromatin Protein 1 (HP1) paralog, HP6/Umbrea arose by duplication of *HP1b* within the *melanogaster* species group, and is absent from the more basal *ananassae* subgroup and the *obscura* outgroup (*1*). The ancestral HP6/Umbrea later lost its chromodomain in the *melanogaster*-subgroup lineage, yielding the CSD-only architecture modeled here (Fig 1A). Importantly, this loss was likely driven by strong, recurrent selection (*1*). As a member of the HP1 gene family, HP6/Umbrea is one of several paralogs that have diversified repeatedly by duplication, each evolving distinct chromosomal distributions and roles (*2–4*). Notably, this family contains paralogs that track actively transcribed chromatin, in addition to HP1’s well-described role in silencing (*5*). The HP1 family exemplifies how lineage-restricted new genes are increasingly recognized as an engine of evolutionary novelty (*6*). Despite the attention HP6/Umbrea has received, the functional consequences of its evolution have remained unclear (*1*, *4*, *7*). Although it is known to bind HP1a and other heterochromatin proteins directly (*8*), a mechanistic and potentially selectable molecular function still remains missing. The aim of this study is to close that gap by providing a candidate, testable function through which HP6/Umbrea could be acted on by selection.

**Figure 1:**
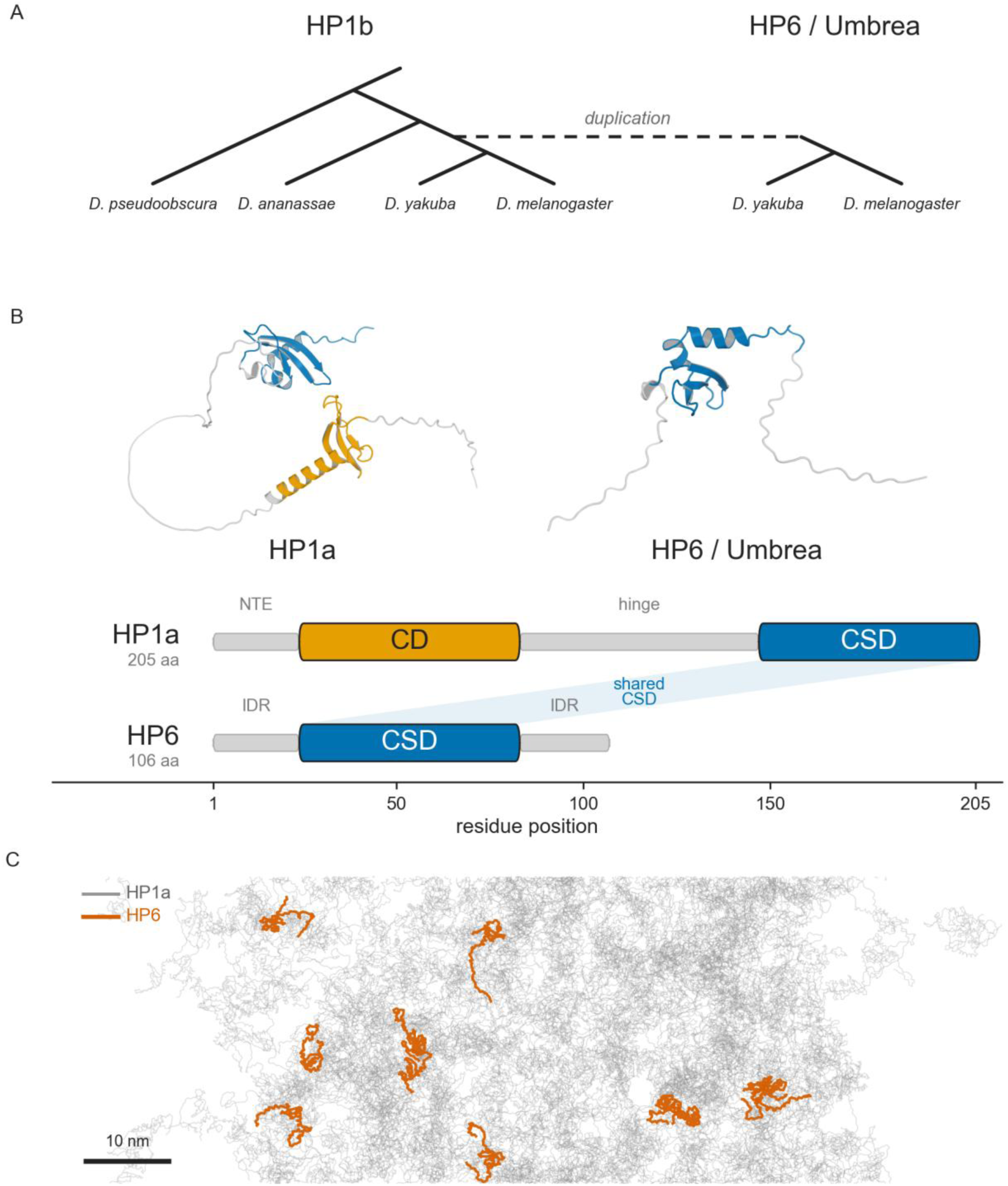
Evolutionary origin and molecular model of HP6/Umbrea. (A) Schematic phylogeny of the HP6/Umbrea origin (branch lengths not to scale): HP6/Umbrea arose by duplication of HP1b in the *melanogaster* species group (absent from *ananassae* and the *obscura* outgroup). While HP1b is HP6/Umbrea’s evolutionary parent, the phase-separating host modeled here is the distinct heterochromatin paralog HP1a (see text). (B) Coarse-grained AlphaFold construct architecture colored by domain (CD orange, CSD blue, disordered gray). HP1a (205 beads) has a chromodomain (CD), disordered hinge, and chromoshadow domain (CSD), while HP6/Umbrea (106 beads) retains only the CSD, flanked by intrinsically disordered regions. (C) Bead-level cross-section of an equilibrated slab. HP1a chains (gray) form the dense condensate, HP6/Umbrea chains (orange) are distributed throughout its interior.

Canonical HP1 proteins are built from three parts: an N-terminal chromodomain that reads the H3K9me mark, a disordered hinge that contacts nucleic acid, and a C-terminal chromoshadow domain (CSD) that mediates homo- and hetero-dimerization (*9–11*). Note that the *Drosophila* HP1 family (HP1a/*Su(var)205*, HP1b, HP1c, HP1d/Rhino, HP1e) diversified largely independently of the mammalian HP1*α*/*β*/*γ* proteins, so the shared letter designations are not statements of orthology. In particular, fly HP1a and HP1b are not the counterparts of human HP1*α* and HP1*β* (*2*, *12*). A leading model for how HP1 builds a heterochromatin compartment is liquid–liquid phase separation (LLPS). Among the fly paralogs, HP1a is the canonical constitutive-heterochromatin protein and the one with directly demonstrated autonomous phase separation. *In vitro*, purified fly HP1a spontaneously demixes into liquid droplets and forms dynamic, fusing foci in *Drosophila* embryos (*13*), whereas the other paralogs have other specialized functions. For example, HP1c requires the zinc-finger proteins WOC and ROW for chromatin binding (*14*), and Rhino and HP1e are germline-restricted (*5*, *12*). We therefore model *HP1a*, not HP6/Umbrea’s parental paralog *HP1b*, as a phase-separating host condensate in our simulations. Note that human HP1*α* likewise forms condensates, but only when it is N-terminally phosphorylated or DNA-bound (*15*). Importantly, the HP1–nucleosome interaction can itself drive condensation. For example, fission-yeast Swi6 (also an HP1-family protein) restructures the nucleosome core to promote phase separation *in vitro* (*16*), and chromatin itself can phase separate in a manner tuned by its modification and binding state (*17*), thus placing heterochromatin within the broader biomolecular-condensate paradigm (*18*, *19*). This model is, however, actively contested. For example, high-resolution imaging in mouse cells finds heterochromatin adopting compacted states without the hallmarks of HP1-driven LLPS (*20*), and whether phase separation is required for heterochromatin compaction *in vivo* remains openly debated (*21*). We therefore adopt LLPS as a working physical framework for the compartment HP6/Umbrea enters, not as an established fact (Discussion). The closest simulation precedent to the present work reproduces the thermodynamics and kinetics of heterochromatin condensate formation from HP1-driven phase separation (*22*), though with a polymer-thermodynamics model rather than the residue-level, sequence-specific representation we use here. Within this framework, a condensate is characterized by a small set of measurable order parameters, obtained here by direct-coexistence “slab” simulation (*23*): many copies of the protein are placed in an elongated box and allowed to demix, separating spontaneously into a protein-rich dense phase, i.e.: the condensate itself forming a box-spanning slab, that coexists at equilibrium with the surrounding protein-poor dilute phase. Importantly, the two phases continually trade molecules across a sharp interface, providing an *in silico* counterpart of a condensate droplet suspended in nucleoplasm. This simulation geometry directly emulates a condensate droplet flattened into a slab so that concentrations can be read cleanly along the long axis. Key parameters then follow by measuring various aspects of the equilibrium concentration profile, notably the saturation concentration *c*_sat_, the dilute-phase level at the phase boundary (below this level the protein remains dissolved; above it, a condensate forms), the dense-phase density *c*_dense_ inside the slab, the interfacial tension *γ* (the energetic cost of the dense–dilute boundary that reflects boundary sharpness), and the partition coefficient *K*_p_ of any client (the ratio of its dense-to dilute-phase concentration). As a result, the model can provide high-quality *in silico* estimates of each quantity, allowing us to evaluate HP6/Umbrea’s proposed function within the model.

HP6/Umbrea’s structure is reduced relative to canonical HP1a (Fig 1B). It retains the CSD (the dimerization module) but has lost the ancestrally shared chromodomain and hinge (*8*, *24*). As a result, HP6/Umbrea possesses no obvious independent chromatin-binding capacity, suggesting that direct chromatin binding is not central to its molecular role *in vivo*. Consistent with this, HP6/Umbrea binds the HP1a protein (*Su(var)205*) directly through a CSD–CSD interaction, and the two co-localize in pericentric heterochromatin and at telomeres *in vivo* (*8*). Because the shared CSD is conserved across the HP1 family, an HP1b-derived CSD can still pair with HP1a. The newly derived HP6/Umbrea has been shown to belong to a heterochromatin-protein network in which HP1 serves as the docking platform (*24*) (Fig 1C) such that its heterochromatic targeting is HP1-dependent. We note, however, that paralog selectivity among the HP1 proteins has not been tested directly. This reduced architecture thus points to a specific role: HP6/Umbrea retains the functional domain that allows HP1 proteins to pair but has lost the domains that bind chromatin and DNA, suggesting that it acts as a modifier rather than an autonomous structural component.

We therefore propose and evaluate a specific candidate function using an *in silico* coarse-grained condensate model (*25–27*). (i) Because HP6/Umbrea keeps only the CSD, its sole route into the HP1a condensate is CSD–CSD dimerization with HP1a, thus acting as an *HP1a-recruited* client, not an autonomous one. (ii) Once inside, HP6/Umbrea acts as a molecular *plasticizer* (*28*, *29*), which loosens the interior packing of the condensate (lowering *c*_dense_) without shifting the phase-separation threshold *c*_sat_, the interfacial tension *γ*, or its host’s partitioning. A generic weakening of cohesion would drop the surface tension too, but a plasticizer (a spacer that dilutes contacts locally without dissolving the phase) is predicted to move the interior density preferentially. (iii) A looser interior is, by standard condensate physics, more permeable to large clients such as RNA polymerase machinery (*28*, *29*), which predicts a position-effect-variegation (PEV)-like modulation of repression, i.e., leaky silencing of otherwise-off heterochromatic genes (*30*). We also highlight that HP1a is the known product of *Su(var)205*, a founding suppressor-of-variegation locus whose loss-of-function relieves silencing, so wild-type HP1a promotes heterochromatic repression (*31*). A condensate-loosening factor that HP1a itself recruits, acting to relax HP1a’s repressive function, would antagonize its own partner’s function from within the heterochromatin condensate. We thus propose that HP6/Umbrea may function in an enhancer-of-variegation-like manner by reducing transcriptional repression via heterochromatin. Because HP6/Umbrea is lineage-restricted and tissue-biased in expression (*7*), its effects may likewise be restricted and tissue-based. This is the type of dose-sensitive, potentially selectable phenotype that a young, rapidly evolving gene could produce (Discussion). Steps (i)–(ii) are addressable in the model, while step (iii) needs to be directly tested. We further ask whether the disordered tail that is implicated in this effect carries a molecular-evolutionary signature consistent with an evolved function.

## Results

### HP6/Umbrea is recruited into the condensate via HP1a

HP6/Umbrea enters the HP1a condensate by riding HP1a, not by partitioning on its own, as its domain architecture predicts. We tested this directly *in silico* using a slab of the fly HP1a/HP1a condensate at 258 K (this is an effective model temperature rather than a direct physical temperature (Discussion)). We then compared four physical placements of the same 106-bead HP6/Umbrea chain: (1–2) covalently tethered in place of one HP1a dimerization arm (an HP1a/HP6 heterodimer, added or substituted), (3) the matched arm-deletion controls, and (4) a free (untethered) folded HP6/Umbrea, each seeded from both a compact and a dispersed configuration.

The partition coefficient of the HP6/Umbrea-bearing chemical species differs clearly by topology (Fig 2). Free HP6/Umbrea barely enters the condensate (ln*K*_p_ = 1.86 ± 0.21, *K*_p_ ≈ 6), whereas tethered HP6/Umbrea partitions approximately sixfold more strongly (ln*K*_p_ = 3.61 ± 0.07 additive, 3.75 ± 0.11 substitutive, *K*_p_ ≈ 37–42). Pooling the two concordant tethered estimates (i.e., compact- and dispersed-seeded) against the free value gives a tether contribution of robust to the additive-vs-substitutive mixing artefact related to *c*_sat_ below. The separation is complete and highly confident, resulting in a +1.82 ± 0.22 shift at roughly an 8*σ* effect, with all four tethered runs at ln*K*_p_ = 3.54–3.86 and both free runs at 1.65–2.07 (no overlap, per-run values in Table S1). Because *K*_p_ is a ratio of the same chemical species across the two phases, this ratio is also robust to the overall-density and probe-count transients that affect the *c*_sat_ endpoints.

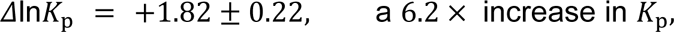

**Figure 2:**
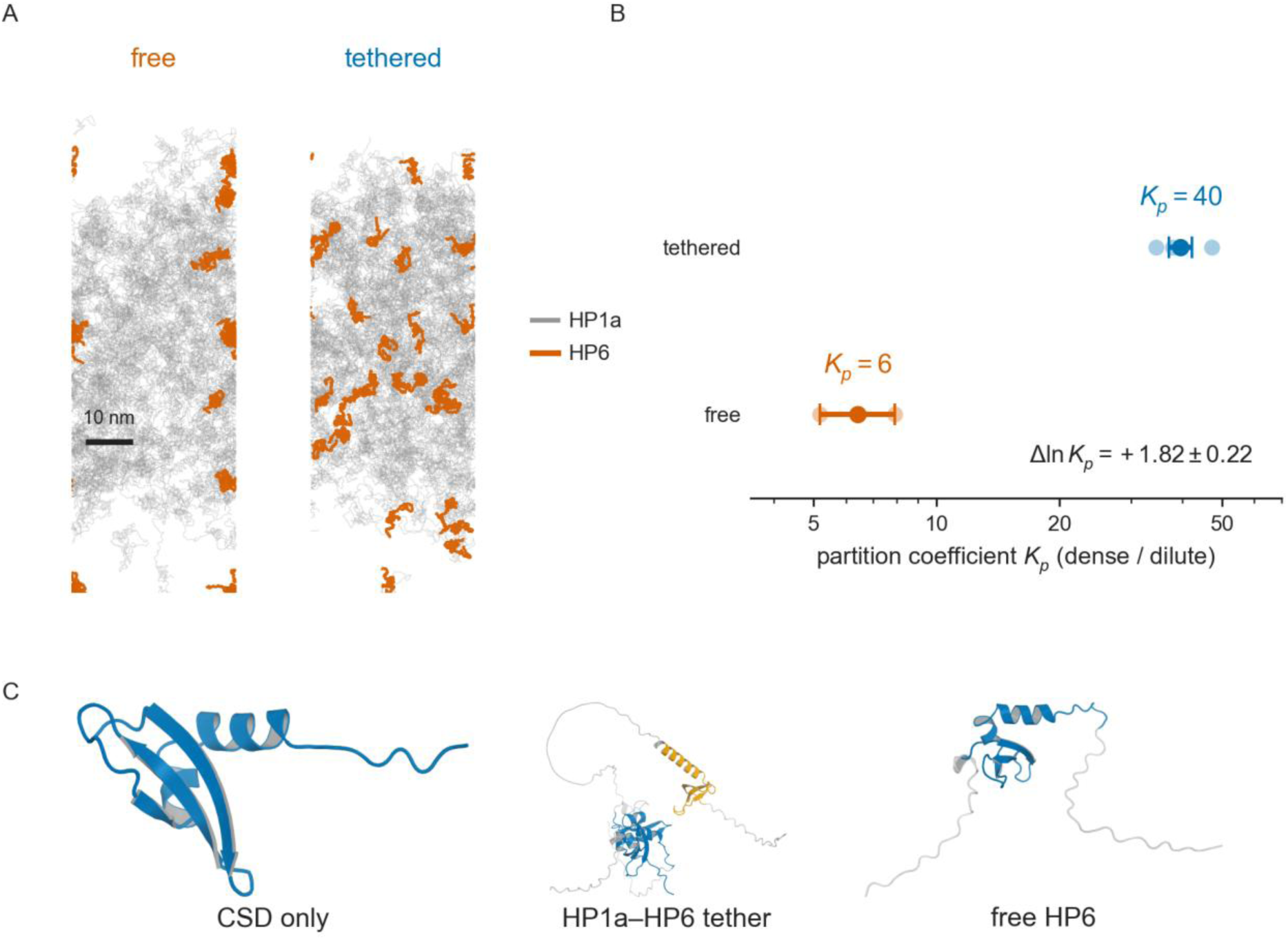
HP6/Umbrea is recruited into the condensate by tethering to HP1a. (A) Bead-level cross-sections of equilibrated frames for free HP6/Umbrea (left) and HP1a-tethered HP6/Umbrea (right) at 258 *K* (*x* = 0.16). HP1a chains are gray, and HP6/Umbrea chains are orange. Free HP6/Umbrea is largely confined to the dilute rim, whereas tethered HP6/Umbrea threads the dense interior. (B) Partition coefficient *K_p_* (dense/dilute) of the HP6/Umbrea-bearing species on a log axis. Shaded markers represent initial conditions (N=4) reflecting compact (N=2) and dispersed (N=2) configurations, whereas solid markers represent bracket midpoint. Tethering raises partitioning from *K_p_* ≈ 6 (free) to *K_p_* ≈ 40 (tethered), a pooled tether contribution of Δln*K_p_* = +1.82 ± 0.22. (C) The three compared constructs as folded cartoons (AlphaFold models, domain-colored as in Fig 1; see Materials and methods). The isolated CSD, the HP1a–HP6/Umbrea tether, and free HP6/Umbrea are shown.

We note that this is the expected sign (i.e., > 0) and the magnitude is consistent with the domain architecture, as a chain that only retains the CSD has little affinity for the condensate alone but is dragged in when covalently coupled to an HP1a particle. This is also consistent with prior observations demonstrating that HP6/Umbrea both localizes to heterochromatin and interacts with HP1a (*24*).

### HP6/Umbrea does not move the threshold, the surface, or the host’s partitioning

If HP6/Umbrea merely destabilized the condensate, rather than acting as a plasticizer, both the phase-separation threshold and the actual interface would be predicted to change. However, three independent condensate order parameters do not vary statistically with HP6/Umbrea dose at 258 K, arguing against a destabilization model. Only the condensate density changed in our simulations (Fig 3A–D). The titration slope *β* = dln*c*_sat_/d*x* over the production dose grid *x* = 0, 0.03, 0.06, and 0.09 shows no HP6/Umbrea-dependent *increase* in the saturation concentration. Depending on the fit, it ranges from indistinguishable from zero on the pooled additive-plus-substitutive set (slope *p* ≈ 0.99) to a small negative value on the cleaner subset (*β* = −3.54 ± 1.29 at *L* = 35 nm). Both estimates are far from, and opposite in sign to, the +8.5 that the MOFF model predicts for pure chain-length dilution. This result does not support HP6/Umbrea-driven destabilization of the phase boundary. Similarly, across *x* ∈ [0,0.09] the tension slope is d*γ*/d*x* = +62 ± 76 *μ*N/m per unit *x* (*p* = 0.43, *n* = 11), with the direct interface-width slope equally null (d*w*/d*x* = −10 ± 10 nm per unit *x*, *p* = 0.37). While tests for amphiphilic heterodimer adsorption at the interface (a surfactant Gibbs excess) are inconsistent across runs, the data consistently yield a null tension slope, providing further support against HP6/Umbrea-driven destabilization. Finally, the partition coefficient of the HP1a/HP1a probe itself does not change with HP6/Umbrea dose (slope null, *p* = 0.70; probe ln*K*_p_ sits at ≈ 4.1–4.3 across all compositions). As the dose changes, HP6/Umbrea can alter its own particle distribution (*HP6/Umbrea is recruited into the condensate via HP1a*) without altering its host’s distribution under simulation conditions.

**Figure 3:**
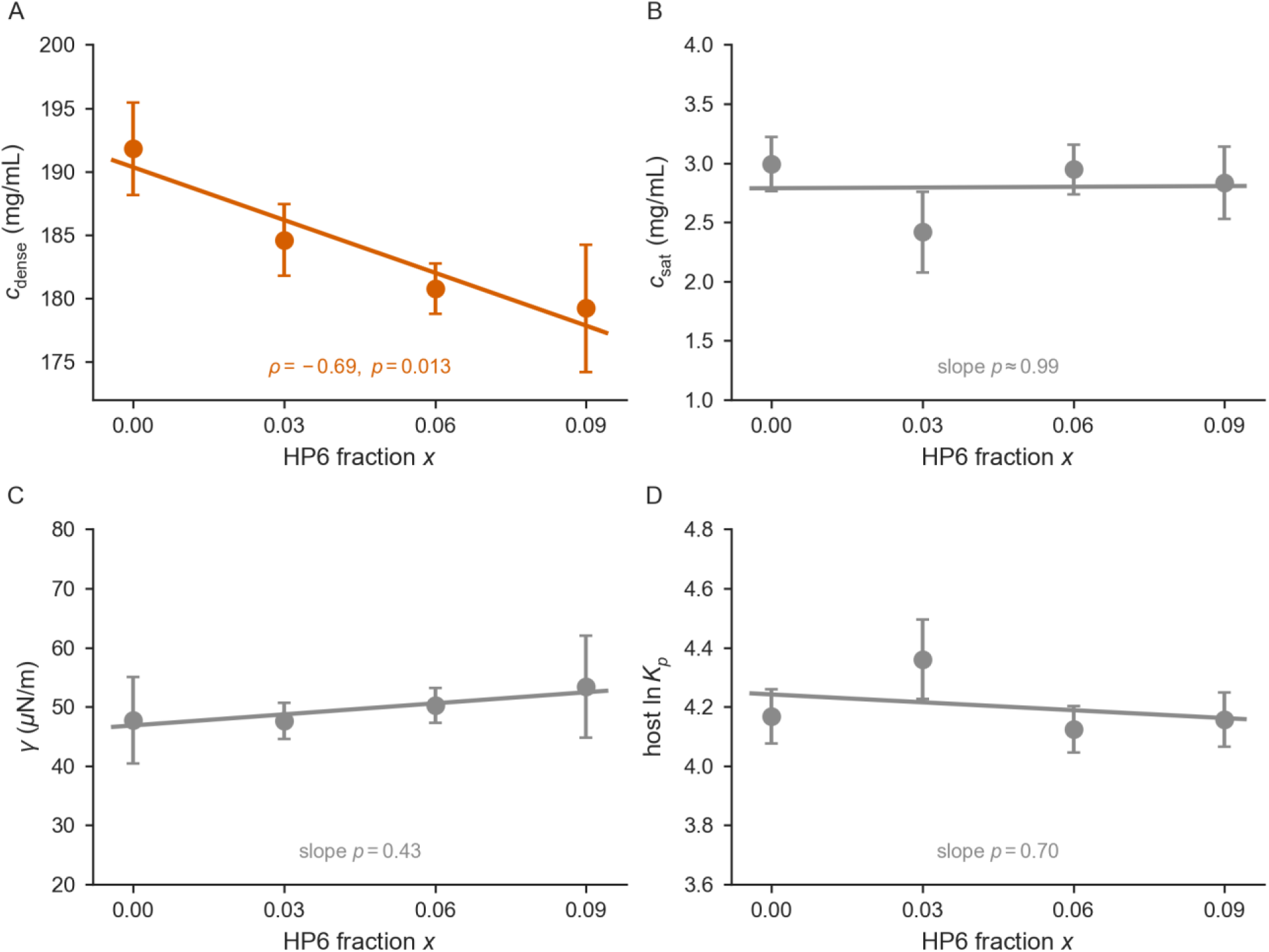
HP6/Umbrea acts as a plasticizer in simulation. Condensate order parameters versus HP6/Umbrea fraction *x* at 258 *K* (*K* = 35 *nm*). Points show per-dose means, indicated by ± standard error (SE) and a regression line. (A) The HP1a host-network dense-core density *c*_*c*_ (orange) falls monotonically (∼7%; Spearman ρ = −0.69, *_p_* = 0.013; total dense-phase density is unchanged, Figure S1). (B) The saturation concentration *c*_*sat*_, (C) the interfacial tension γ, and (D) the host HP1a/HP1a partition coefficient each remain statistically flat (slope *_p_* ≈ 0.99, 0.43, and 0.70, respectively). The specific response of *c*_*c*_ to HP6/Umbrea dose argues against a generic loss of cohesion.

### HP6/Umbrea loosens the dense-phase interior

In contrast to these parameters, one measured order parameter does change: the HP1a host-network density in the dense core, *c*_dense_, falls monotonically as the HP6/Umbrea fraction rises (Fig 3A; per-dose values in Table S2). While the *total* dense-phase density is essentially unchanged across dose (added HP6/Umbrea substitutes into the volume vacated by HP1a; Figure S1), the density of the cohesive HP1a–HP1a network decreases. Under simulated conditions, the overall mass of the network stays roughly constant, indicating a simple expansion of condensate volume. Over the accessible range, the decrease in density is measured at ∼7%, a moderate but statistically significant change. The rank correlation between *c*_dense_ and HP6/Umbrea fraction yields Spearman *ρ* = −0.69 (*p* = 0.013, *n* = 12 replicates) and is corroborated by a permutation test on the slope (*p* ≈ 0.02, Pearson *r* = −0.67). Notably, the correlation analysis pools all completed replicates, including three with low *c*_sat_ exchange counts, since the dense-core density needs no interfacial exchange and is converged by ∼1 μs (Figure S2).

The pattern observed across the examined order parameters, rather than any single value, helps distinguish among possible mechanisms for HP6/Umbrea’s effects. Importantly, three possible explanations here predict different signatures. A generic loss of cohesion (weaker inter-chain attraction) is expected to lower the interfacial tension and raise *c*_sat_ as it lowers *c*_dense_. Instead, only *c*_dense_ moves while *γ*, *c*_sat_, and host *K*_p_ stay flat, thus arguing against a generic loss of cohesion. Two possibilities remain that could produce the same *c*_dense_-specific signature: (1) Generic chain-length dilution, in which any shorter CSD-bearing chain would displace HP1a and lower the host-network density, or (2) an HP6/Umbrea-specific plasticizer effect. However, simulation alone cannot distinguish them. We therefore examined molecular-evolutionary evidence to ask whether HP6/Umbrea’s sequence is functionally distinct.

### Comparative sequence analysis highlights a previously under appreciated C-terminal tail

Across the *melanogaster* group, the chromoshadow domain is constrained (nonsynonymous-to-synonymous rate ratio dN/dS = 0.285), whereas its flanking disordered tails evolve roughly twofold faster (dN/dS = 0.503). Notably, these disordered tails overlap the exact regions implicated by our model in the interior-loosening effect (Materials and methods). The tails are thus the least-constrained portion of the HP6/Umbrea protein, coinciding with its most recent, functionally important innovations that confer species-specific centromere localization (*1*). As dN/dS estimates from ∼47 tail codons cannot separate relaxed constraint from positive selection, we therefore applied PAML codon models (*Materials and Methods*) across the *melanogaster* group and three chromodomain-retaining outgroup lineages, with the branch to the *melanogaster*-subgroup ancestor as the foreground. Surprisingly, no domain shows significant positive selection on that branch (branch-site model A: chromoshadow domain 2ΔlnL = 3.2, C-terminal tail 2ΔlnL = 2.1, both non-significant). This may partially reflect that the disordered C-terminal tail is *largely absent* from the pre-neofunctionalization outgroups. Like the loss of the chromodomain, its origin appears to be an architectural innovation of the *melanogaster* subgroup, not a codon-level selective sweep. Within the subgroup, however, the tail itself evolves under some level of purifying selection (dN/dS ≈ 0.5). We note that this rate of evolution is even looser than the chromoshadow domain (dN/dS ≈ 0.3) but well below neutrality. The C-terminal tail therefore carries the signature of a young, functionally constrained module, consistent with an evolved role. We emphasize, however, that these rate-based tests cannot attribute this constraint to the condensate-plasticizing function specifically, rather than to centromere targeting.

### Internal loosening as a packing effect, not a short-time mobility increase

To determine the effect of HP6/Umbrea on the internal dynamics of the simulated condensate, we measured the lateral self-diffusion of the HP1a/HP1a probe inside the dense core using the same trajectories. Interestingly, the interior mobility of HP1a/HP1a homodimers does not rise with HP6/Umbrea: the measured slope is d*D*/d*x* = −0.256 ± 0.596 nm^2^/ns per unit *x* (*p* = 0.68; Spearman *ρ* = −0.09, *p* = 0.79; Fig 4A). As an internal control, the dense phase is, as required, far slower than the dilute phase (*D*_dense_ ≈ 1.91 vs *D*_dilute_ ≈ 66 nm^2^/ns, a factor ∼35; Fig 4B), confirming that the measurement resolves the two environments.

**Figure 4:**
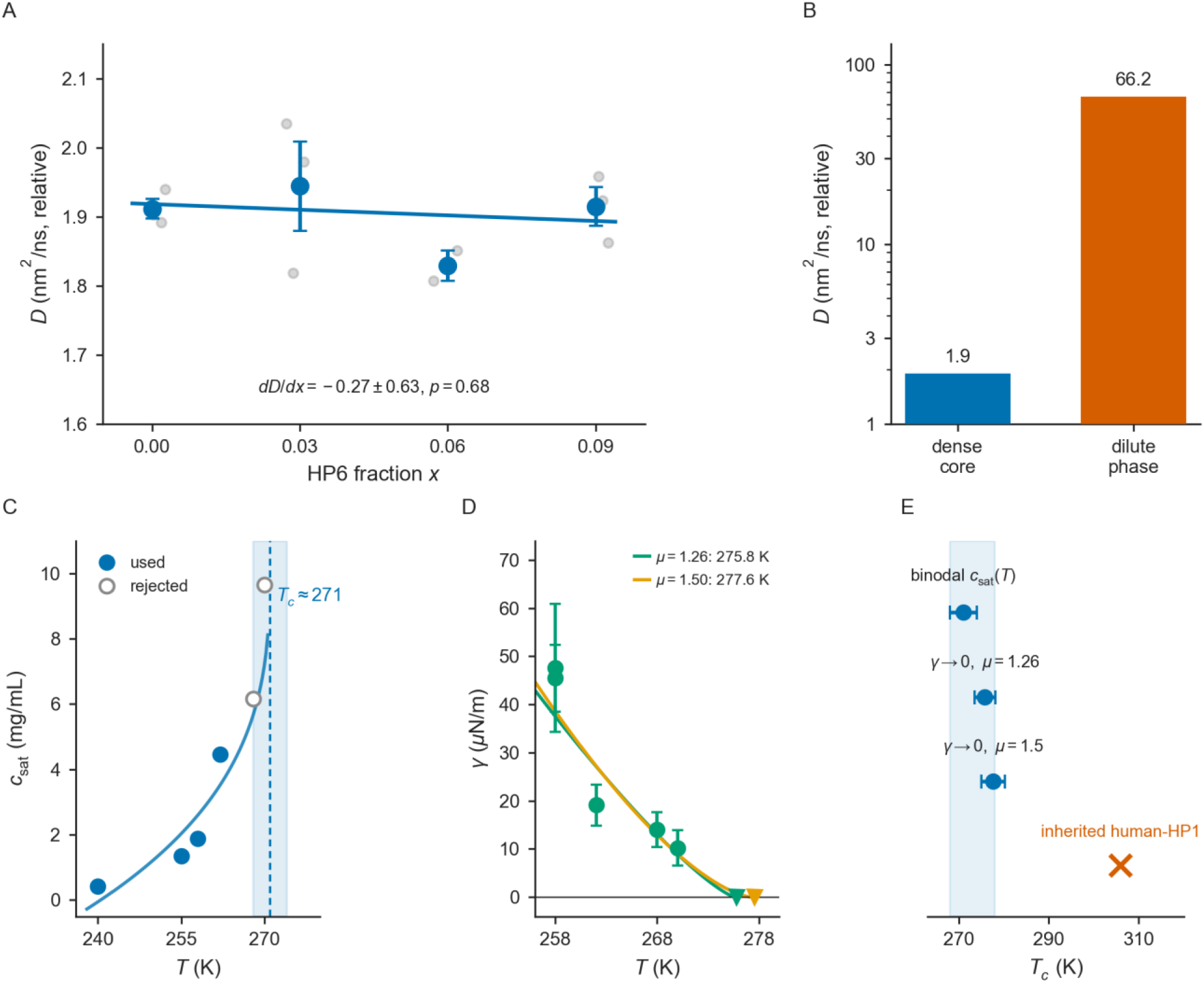
Internal dense-phase dynamics and the *Drosophila* HP1a critical temperature. (A) Lateral self-diffusion *D* of the HP1a probe in the dense core versus HP6/Umbrea fraction *x* (means ± SE). Measured slope across conditions is null (*dD*/*dx* = −0.26 ± 0.60 *nm*^2^/*ns*, *_p_* = 0.68), consistent with a density/packing effect, not a short-time mobility increase. (B) Dense-core diffusion is ∼35× slower than the dilute phase (*D_dense_* ≈ 1.9 vs *D_dilute_* ≈ 66), confirming the probe resolves the two environments. Note that *D* is a relative, fictitious-time coarse-grained coefficient compared across doses only. (C) Dilute-branch *c*_*sat*_(*T*) extrapolation places *Tc* ≈ 271 *K* (shaded range [268,274] K, open points above ∼270 K excluded from the fit owing to recurrent spontaneous aggregation, see text). (D) Fits of γ(*T*) = γ_0_(*Tc* − *T*)^μ^ yield *Tc* = 275.8 ± 2.3 K (3D-Ising, μ = 1.26) and 277.6 ± 2.6 K (mean-field, μ = 1.50). (E) A binodal estimate (≈271 K) and the two interfacial estimates all depart from the previously simulated human-HP1 value near 306 *K* (shaded band, adopted range).

Notably, the interior loosening reported above manifests as reduced packing density, not as increased short-time lateral mobility. We therefore frame the proposed mechanism as a spacer/packing effect rather than a change in mobility or viscosity. Importantly, the coarse-grained *D* is a relative, fictitious-time mobility measure that is meaningful only across doses at a fixed model, and our probe reports a short-time, lateral, mildly subdiffusive (*α* ≈ 0.9) coefficient that may not track a true long-time viscosity. The step from “looser packing” to “more permeable to large machinery” is thus a physical *inference* from reduced density (*29*), which a longer-time or absolute-mobility probe could more directly measure.

### The Drosophila HP1a critical temperature

Earlier project stages assumed a critical temperature carried over from human HP1. Because a coarse-grained model’s transition temperature is calibration-dependent (Discussion, *Simulated temperatures*), we estimate the *Drosophila* HP1a critical temperature *T*_c_ directly, by two independent routes: 1) the difference between dense and dilute phases, and 2) directly on the interface between dense and dilute phases Fig 4C–E). The binodal route extrapolates the coexisting densities and places *T*_c_ ≈ 271 K (adopted range [268,274] K; 68% bootstrap [265.7,271.0] K). An orthogonal, interfacial route utilizes the fact that *γ* → 0 at the critical point. Thus, fitting *γ*(*T*) = *γ*_0_(*T*_c_ − *T*)*^μ^* estimates *T*_c_ via the interface between the dense and dilute phases, independently of coexisting density measurements. This yields *T*_c_ = 275.8 ± 2.3 K for the 3D-Ising exponent *μ* = 1.26 and 277.6 ± 2.6 K for mean-field *μ* = 1.50.

While these estimates are orthogonal and utilized differing simulation features, the two estimates are not independent readings of the same number, as the binodal methodology places *T*_c_ at 270–272 K, while the interfacial methodology estimates a value that is ∼4–5 K higher. This offset is expected by theory, as the single-box tension is a *lower* bound on *γ*, thus biasing the extrapolated *T*_c_ upward. We therefore take the binodal (≈ 271 K) as the preferred estimate and the interfacial value as an upward-biased validation measurement. Regardless, both are significantly lower than the previously simulated human-HP1 value (∼306 K under the same model (*25–27*)). A third, separate, model-free indicator is consistent with this value, as the simulated slab spontaneously pinches into droplets above ∼270 K (96% of frames split at 272 K), thus placing *T*_c_ near 270–272 K.

## Discussion

### A candidate evolutionary–biochemical function

The evolutionary history of HP6/Umbrea, including its rapid sequence divergence (*1*, *4*) and its regulatory turnover (*7*), is documented. Despite previous work, a full molecular function has remained elusive. The central contribution of this study is a candidate mechanism for HP6/Umbrea’s molecular function *in vivo*. HP6/Umbrea’s reduced architecture (CSD retained, chromodomain and hinge lost) predicts, and the simulations reproduce, that it cannot enter heterochromatin autonomously and must be recruited by CSD–CSD dimerization with HP1a (*HP6/Umbrea is recruited into the condensate via HP1a*). Once inside, it acts as a plasticizer, loosening the interior packing while leaving the phase boundary, the surface, and the host’s partitioning intact (*HP6/Umbrea does not move the threshold, the surface, or the host’s partitioning*, *HP6/Umbrea loosens the dense-phase interior*). Importantly, it does not dissolve the phase as a competitor would. This defines an architecture-consistent biochemical role in which HP6/Umbrea is a conditional, HP1a-gated modifier of condensate interior state. In terms of condensate composition, HP6/Umbrea is a scaffold-recruited client that tunes the material state of the phase rather than its existence (*32*). Alternatively, under the stickers-and-spacers model, it contributes spacer-like volume that locally dilutes contacts without adding the cohesive stickers that would move the phase boundary (*33*).

### A prediction for repression: a PEV-like rheostat

Beyond the broad interest in heterochromatin biochemistry, we aimed to understand the phenotypic effects through which natural selection could act on HP6/Umbrea. One predicted consequence of our findings relates to gene regulation. Condensates with looser, lower-density interiors are expected to be more permeable to large client molecules (*29*). Such clients would include the transcription and remodeling machinery whose *exclusion* helps enforce the silencing function of heterochromatin. The observation that condensates can, in other systems, gate the access of transcriptional machinery and thereby shape gene control (as at coactivator-condensed super-enhancers (*34*)) offers a key analogy for how a permeability change of heterochromatin might couple to transcription. An HP1a-gated interior softener would thus suggest a role for HP6/Umbrea in producing a position-effect-variegation-like modulation of transcription: dynamically tuning the leaky silencing of otherwise-off heterochromatic genes (*30*). We therefore propose HP6/Umbrea as a candidate repression *rheostat* rather than an on/off switch, fitting the phenomenology of a fast-evolving, dose-sensitive heterochromatin modifier.

Under such a model, these results give the rheostat a specific genetic identity. HP1a is encoded by *Su(var)205* (*Su(var)2-5*), one of the founding *suppressor-of-variegation* loci. Reducing its dose suppresses PEV, so wild-type HP1a is required to build and maintain repressive heterochromatin (*30*, *31*). PEV is known to be a graded, strongly dose-dependent readout in which modifier loci act by mass action, with haplo-suppressor/triplo-enhancer dosage series (*30*, *35*). Thus, a factor that changes the molecule count or relative degree of packing of heterochromatin components is expected to move silencing *quantitatively* rather than as a binary switch. Our model places HP6/Umbrea on the opposite side of that balance within the same machinery. Here, HP6/Umbrea is recruited by HP1a (an interaction demonstrated *in vivo* (*8*)), thus loosening the interior. This may relax silencing in the same direction as reducing (functional) HP1a. In classical genetic terms, HP6/Umbrea would produce an enhancer-of-variegation-like phenotype, de-repressing activity imposed by the Su(var) protein it partners with. Specifically, it is an HP1a-recruited brake on HP1a’s own silencing. HP6/Umbrea is a recently evolved, lineage-restricted paralog with germline-biased, rapidly evolving expression (*2*, *4*, *7*) that is dispensable for viability and fertility (*4*). We note that these observations are compatible with a tissue-biased modulator rather than a role as a central component of the heterochromatin machinery. While this mirrors the observation that another HP1 paralog can alter a host condensate’s material state (mammalian HP1*β* dissolves HP1*α* condensates (*36*)), we are unaware of a prior proposal that an HP1 paralog tunes PEV as an HP1a-recruited de-repressing rheostat.

The tissue in which HP6/Umbrea is most abundant points to where such an activity would act. HP6/Umbrea’s expression is tissue- and testis-biased (*7*), and the *Drosophila* testis is well known for transcribing an unusually large and complex fraction of the genome. This includes many lineage-specific, *de novo*, and otherwise-silenced sequences (*37*, *38*). Such a permissive, “leaky” program has been linked to a globally open germline chromatin state and to the emergence of new genes (*39–41*). Notably, a recent study has demonstrated that the open chromatin that permits the expression of still-segregating *de novo* genes is itself a derived state, suggesting that HP6/Umbrea’s enhancer-of-variegation function may be a key step in exposing these young open reading frames first to the cell’s transcriptional and translational machinery, and consequently to natural selection (*42*). However, most transcriptional leakiness is broad and largely euchromatic suggesting that heterochromatin de-repression may be only one component of many in driving leakiness. We therefore propose that HP6/Umbrea may be one of several factors that help open otherwise-silenced chromatin in the male germline, placing a young, rapidly evolving heterochromatin paralog among the contributors to one of the least-understood features of the testis transcriptome. Consistent with a *modulatory* rather than obligatory role, HP6/Umbrea has been found to be dispensable for viability and fertility under laboratory conditions (*4*). A dispensable gene can nonetheless be under selection is not a contradiction, since the evidence for selection is its recurrent, rapid diversification (*1*), not a knockout phenotype. We stress, however, that our coarse-grained HP1a/HP6 condensate contains no DNA, no nucleosomes, and no RNA polymerase, leaving open the possibility that HP6/Umbrea may not act as a plasticizer *in vivo*.

### Simulated temperatures

We note that the absolute temperature values here, with a production point of 258 K and a fly critical temperature near 271 K, are *effective* temperatures of a transferable coarse-grained force field, decoupled from physical readings that map one-to-one onto degrees in a nucleus. MOFF’s energy scale is fixed by the experimental phase behavior it was parameterized against (*25*, *26*), so its transition temperatures inherit that calibration. For example, when simulations run with the earlier human-HP1–derived constants, the same model places the critical point near 306 K. Re-anchoring of simulation parameters to the *Drosophila* protein shifts this critical point to 271 K (*The Drosophila HP1a critical temperature*). That this value tracks with the reference is critical, as the model may reliably resolve relative temperature changes rather than absolute temperature in Kelvin (working point *T*/*T*_c_ ≈ 0.95). Importantly, the ordering of states and the sign and size of shifts are all evaluated *at a single temperature*. All comparisons in this study were made at one fixed 258 K such that the effective-temperature scale cancels from the HP6/Umbrea effect entirely.

## Materials and methods

### Coarse-grained model

Proteins were represented with the MOFF one-bead-per-residue coarse-grained force field (*25*), which reproduces homolog-resolved HP1 phase behavior (*26*), as implemented in OpenABC (*27*). MOFF sits within a family of residue-level, sequence-dependent coarse-grained models developed for biomolecular phase separation (*43–45*). We adopt it here for its calibrated, one-bead-per-residue treatment of HP1, in preference to the more finely grained near-atomistic alternatives (e.g., Martini-based and adapted force fields) whose per-residue cost is impractical for the many-replica titration performed here (*46*, *47*). OpenABC also provides the MRG coarse-grained double-stranded DNA model (*48*). We note that the HP1a/HP6 condensates studied here are protein-only (no DNA). Each fly HP1a chain is 205 beads with a chromodomain (residues 24–82), a disordered hinge (83–146), and a chromoshadow domain (CSD, 147–205), with the chromodomain and CSD held folded by native pairs and dimerization mediated by the CSD. HP6/Umbrea is a 106-bead chain that comprises an N-terminal intrinsically disordered region (IDR; 1–23), the CSD (24–82), and a C-terminal IDR (83–106), with only the CSD folded and, again, CSD-mediated dimerization, carrying neither a chromodomain nor a hinge. The folded domains and their intra- and inter-chain native contacts were defined from AlphaFold structural models (the HP1a monomer from the AlphaFold Protein Structure Database, accession AF-P05205, and the HP6/Umbrea monomer and the HP1a–HP6/Umbrea assembly from AlphaFold predictions). An HP1a/HP6 heterodimer forms a CSD–CSD interface with 47 inter-chain native pairs, compared with 52 for an HP1a/HP1a homodimer.

### Slab direct-coexistence simulations

Phase behavior was obtained by the slab direct-coexistence protocol (*23*). A condensed slab spanning the periodic box in *x*, *y* is elongated along *z*, and the coexisting dense and dilute concentrations are read from the equilibrated density profile. Production runs used a 35 × 35 × 400 nm box with 200 HP1a/HP1a homodimers and an HP6/Umbrea dose grid *x* = 0, 0.03, 0.06, and 0.09 (three replicates per dose). The recruitment (tether) tier used six compositions at *x* = 0.16, each from a compact and a dispersed seed. Exploratory temperature and box-size tiers used a 25 nm lateral box. Dynamics were integrated with a Langevin thermostat (collision frequency 1 ps^−1^) at a 5 fs timestep, with configurations saved every 0.5 ns. Production length was 3.25 μs per run at 258 K (PROD_US), set by the slowest-exchanging arm. The primary salt concentration was 82 mM, with 100 mM used as a forking-path guard. The production temperature, 258 K, is 0.95 *T*_c_, the highest temperature at which the perturbed (HP6/Umbrea-bearing) arms retain slab integrity, since HP6/Umbrea is interfacially active and lowers the pinching temperature. As in any transferable coarse-grained force field, this is an effective model temperature set by MOFF’s parameterization (*25*). We therefore report it as a fraction of the model’s own critical temperature (0.95 *T*_c_) and treat absolute values as calibration-dependent.

### Endpoints

For each run we computed the saturation concentration *c*_sat_ as the dilute-phase HP1a/HP1a concentration (using the split-frame–filtered value where slab pinching contaminated frames), the dense-phase density *c*_dense_ as the HP1a/HP1a concentration in the dense core |*z*| ≤ 10 nm, the partition coefficient *K*_p_ of each species as its dense/dilute ratio, and the interfacial tension *γ* from capillary-wave broadening of the interface width (*49*) using per-molecule center-of-mass *z* (a single-box estimator returns a lower bound *γ* ≥ *c k_B_T*ln(*L*/*B*)/2*πw*^2^). The HP6/Umbrea titration endpoint is the slope *β* = dln*c*_sat_/d*x*, and the critical temperature was estimated both by extrapolating the binodal and by fitting *γ*(*T*) = *γ*_0_(*T*_c_ − *T*)*^μ^* to its zero crossing. Internal self-diffusion *D* was computed from the lateral mean-squared displacement of in-band molecules (MSD*_xy_* = 4*Dτ*, *τ* ∈ [1,5] ns). *D* is a relative, fictitious-time coefficient compared across doses only.

### Uncertainty and convergence

Uncertainties follow a two-level model. *Within* a run the precision of ln*c*_sat_ is set by interfacial exchange rather than by standing headcount, 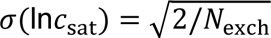 and runs were gated at *N*_exch_ ≥ 250 (i.e. *σ* = 0.089) evaluated on the plateau exchange count, not the naive per-frame count, which understates the true error by < 10 ×. *Between* runs, replicate spread within a dose sets the pooled error, and in the tether tier the compact/dispersed seed pair defines a two-sided convergence bracket whose half-width is the reported uncertainty. On the production dose grid, 8 of 11 completed replicates pass the plateau gate, with every dose retaining ≥ 2 passing replicates so the within-dose error model survives. Three short replicates suffered late equilibration discards (0.52–0.98 μs of production) rather than counting artefacts. At 258 K the 35 nm box is adequate for the *slope* endpoint (the composition-dependent, slope-corrupting part of the finite-size bias is unresolved *Δβ* = −0.71 ± 1.29, 0.5*σ*). We note, however, that absolute *c*_sat_ values were not box-converged (∼0.6–0.7 ln-unit box dependence), requiring further re-checking at temperatures closer to *T*_c_. The *c*_dense_ endpoint converged rapidly, showing no formation transients in saved data. ∼1 μs of simulation averaging converges to ∼2%, requiring neither long equilibration nor the 3.25 μs exchange gate that *c*_sat_ does.

### Sequence divergence and selection analysis

HP6/Umbrea coding sequences were retrieved from NCBI for eight *melanogaster*-group species (*D. melanogaster* NM_134998; *D. simulans* KMY88099; *D. mauritiana* AGG11727; *D. yakuba* EDW87653; *D. teissieri* AGG11725; *D. santomea* AGG11724; *D. erecta* EDV57748; *D. orena* AGG11726) and further outgroup species that have retained their chromodomain in HP6/Umbrea (*D. takahashii* AGG11730; *D. elegans* AGG11735; *D. ficusphila* AGG11738). Each sequence was verified in open reading frame, and the *D. melanogaster* sequence translated exactly to the 106-residue reference protein. Because the basal outgroups retain the ancestral N-terminal chromodomain and hinge that were lost along the *melanogaster*-subgroup stem, the two retaining the chromodomain (*D. elegans*, trimmed to residues 92–154; *D. ficusphila*, to 79–167) were cut at the conserved chromoshadow-domain start so that only the homologous chromoshadow-domain-and-tail region was compared. Protein sequences were aligned with MAFFT (L-INS-i)(*50*) and codons were threaded back onto the protein alignment.

Domains were defined in the *D. melanogaster* coordinate frame: the chromoshadow domain (CSD, residues 24-82) and the flanking disordered tails (N-terminal, residues 1-23, C-terminal residues 83-106). Nonsynonymous and synonymous substitution rates (dN, dS) and their ratio were estimated within the *melanogaster* subgroup by the Nei–Gojobori counting method (*51*) with Jukes–Cantor correction. Domain differences in the nonsynonymous:synonymous count ratio were assessed by a one-sided Fisher exact test. Before fitting codon models, we screened for synonymous-site saturation with pairwise dS (yn00)(*52*).

Maximum-likelihood codon models were fitted per domain in PAML (codeml 4.10.10)(*53*) on the *melanogaster*-group species tree (*54*), with codon frequencies from the F3×4 model, a transition/transversion ratio estimated from data, the universal genetic code employed, and no removal of gapped columns. Three families of models were used. Site models on the unlabeled tree tested for among-site variation in dN/dS and for positively selected sites (one-ratio for M0, nearly-neutral vs positive-selection pair M1a/M2a, and beta vs beta-plus-dN/dS pair M7/M8). A two-ratio branch model then assigned a separate dN/dS to the branch leading to the *melanogaster*-subgroup common ancestor (stem lineage on which derived architecture arose) vs. the remainder of the tree. Finally, a branch-site model A (*55*) tested for episodic positive selection affecting a subset of codons on the foreground branch, comparing the alternative model (foreground (dN/dS)_2_ allowed to exceed 1) against the constrained null ((dN/dS)_2_ fixed at 1). As the N-terminal region is not homologous across the neofunctionalization event, these clade-spanning codon models were fitted for the CSD and the C-terminal tail only. Nested models were compared by likelihood-ratio tests against a χ² distribution (df = 2 for M2a/M1a and M8/M7; df = 1 for the branch and branch-site tests), with the branch-site test evaluated against the recommended 50:50 mixture of χ²₀ and χ²₁. Bayes Empirical Bayes posterior probabilities were used to identify candidate selected sites only where the corresponding likelihood-ratio test was significant.

### Data and code provenance

All quantitative values were read from the collected per-run summaries. Code release and archived inputs are given in the Data Availability statement below.

### Use of generative AI

The authors used a generative artificial-intelligence assistant, Claude Opus 4.8 (Anthropic), in preparing this study. It was used to help write and debug the analysis and figure-generation code, to orchestrate and monitor the coarse-grained simulation campaign on rented compute, to search the literature, and to edit the manuscript. All AI-assisted outputs were checked by the authors: code was inspected and its numerical results independently re-derived where feasible, every reported statistic and figure was verified against the underlying simulation data, and all citations were confirmed against the primary sources. The tool is not an author and generated none of the study’s data (all data derive from the simulations described above). The authors take full responsibility for the content and conclusions.

## Acknowledgments

U.L. would like to thank Yoonji Kim for helpful discussions. The authors would like to thank the Rockefeller University High-Performance Computing Core Facility (RRID: SCR_025889) for their assistance.

## Author contributions

**UnJin Lee (UL):** Conceptualization, Data curation, Formal analysis, Funding acquisition, Investigation, Methodology, Project administration, Resources, Software, Supervision, Validation, Visualization, Writing – original draft, Writing – review & editing. **Li Zhao (LZ):** Funding acquisition, Resources, Supervision, Writing — review & editing.

## Data availability

All analysis and simulation code and the derived data supporting the findings are available at https://github.com/LiZhaoLab/md_hp6, to be made public and archived with a Zenodo DOI upon acceptance. The deposit comprises the simulation inputs, the per-run summary arrays (com_z.npy, com_xy.npy, csat.json, density_profile.csv), the derived result documents (tether_topology_v1, interfacial_tension_v1, working_point_dTc_v1, convergence_boxsize_qc_v1, internal_diffusion_v1) with their machine-readable JSON, and all analysis and simulation code (built on OpenABC (*27*)).

## Supporting information

**Table S1.** Per-run endpoint summary. *c*_sat_, *c*_dense_, the single-box *γ* bound, and per-species ln*K*_p_ for every completed prod_modeA replicate and every tier0_tether run at 258 K, with plateau exchange counts and the *N*_exch_ ≥ 250 pass/fail gate.

| Run | $c_{\text{sat}}$ | $c_{\text{dense}}$ | $\gamma$ | host | 2nd | $N_{\text{exch}}^{\text{plat}}$ | gate |
| --- | --- | --- | --- | --- | --- | --- | --- |
| | mg/mL | mg/mL | $\mu\text{N/m}$ | $\ln K_p$ | $\ln K_p$ | | $\geq 250$ |
| 0.00/r0 | 2.54 | 197.2 | 38.8 | 4.35 | – | 275 |  |
| 0.00/r1 | 3.29 | 193.4 | 42.1 | 4.07 | – | 288 |  |
| 0.00/r2 | 3.14 | 184.8 | 62.2 | 4.08 | – | 106 | × |
| 0.03/r0 | 1.78 | 179.8 | 53.7 | 4.62 | 3.46 | 129 | × |
| 0.03/r1 | 2.94 | 189.6 | 43.4 | 4.17 | 2.37 | 287 |  |
| 0.03/r2 | 2.53 | 184.4 | 45.7 | 4.29 | 3.54 | 277 |  |
| 0.06/r0 | 3.22 | 182.6 | 55.6 | 4.04 | 4.59 | 369 |  |
| 0.06/r1 | 3.09 | 176.8 | 45.4 | 4.05 | 3.68 | 279 |  |
| 0.06/r2 | 2.53 | 182.8 | 49.6 | 4.28 | 3.80 | 258 |  |
| 0.09/r0 | 2.31 | 176.7 | 62.0 | 4.33 | 4.18 | 101 | × |
| 0.09/r1 | 2.79 | 172.1 | 44.7 | 4.12 | 3.78 | 298 |  |
| 0.09/r2 | 3.38 | 188.9 | 38.0 | 4.02 | 2.99 | 301 |  |
| base/com | 3.52 | 189.2 | – | 3.98 | – | 267 |  |
| base/dis | 2.90 | 184.1 | – | 4.15 | – | 487 |  |
| sub_del/com | 2.24 | 178.2 | – | 4.38 | 3.51 | 363 |  |
| sub_del/dis | 2.61 | 170.9 | – | 4.18 | 3.33 | 260 |  |
| sub_teth/com | 3.20 | 171.1 | – | 3.98 | 3.85 | 387 |  |
| sub_teth/dis | 2.45 | 159.2 | – | 4.17 | 3.63 | 440 |  |
| add_del/com | 2.74 | 179.7 | – | 4.18 | 3.41 | 405 |  |
| add_del/dis | 2.76 | 174.7 | – | 4.15 | 3.48 | 453 |  |
| add_teth/com | 2.59 | 172.0 | – | 4.20 | 3.54 | 12 | × |
| add_teth/dis | 2.25 | 172.9 | – | 4.34 | 3.68 | 364 |  |
| free/com | 2.20 | 187.3 | – | 4.44 | 1.65 | 335 |  |
| free/dis | 2.67 | 183.2 | – | 4.23 | 2.07 | 253 |  |

**Table S2.** Condensate order parameters vs HP6/Umbrea fraction. Per-dose values underlying Fig 3. At 258 K (*L* = 35 nm, prod_modeA), *c*_dense_ is the pooled per-dose mean of the dense-core density (|*z*| ≤ 10 nm) over the completed replicates. *c*_sat_ slope, *γ* slope, and host-*K*_p_ slope are all statistically null (*HP6/Umbrea loosens the dense-phase interior*). *indicates that one replicate did not pass QC due to shifting of slab center of mass.

| $x$ | reps | $c_{\text{dense}}$<br>(mg/mL) | $\gamma$ ($\mu\text{N/m}$) |
| --- | --- | --- | --- |
| 0.00 | 3 | $191.8 \pm 3.6$ | $47.7 \pm 7.3$ |
| 0.03 | 3 | $184.6 \pm 2.8$ | $47.6 \pm 3.1$ |
| 0.06 | 3 | $180.7 \pm 2.0$ | $50.2 \pm 3.0$ |
| 0.09 | 3* | $179.2 \pm 5.0$ | $53.4 \pm 8.7^*$ |

**Fig S1.**
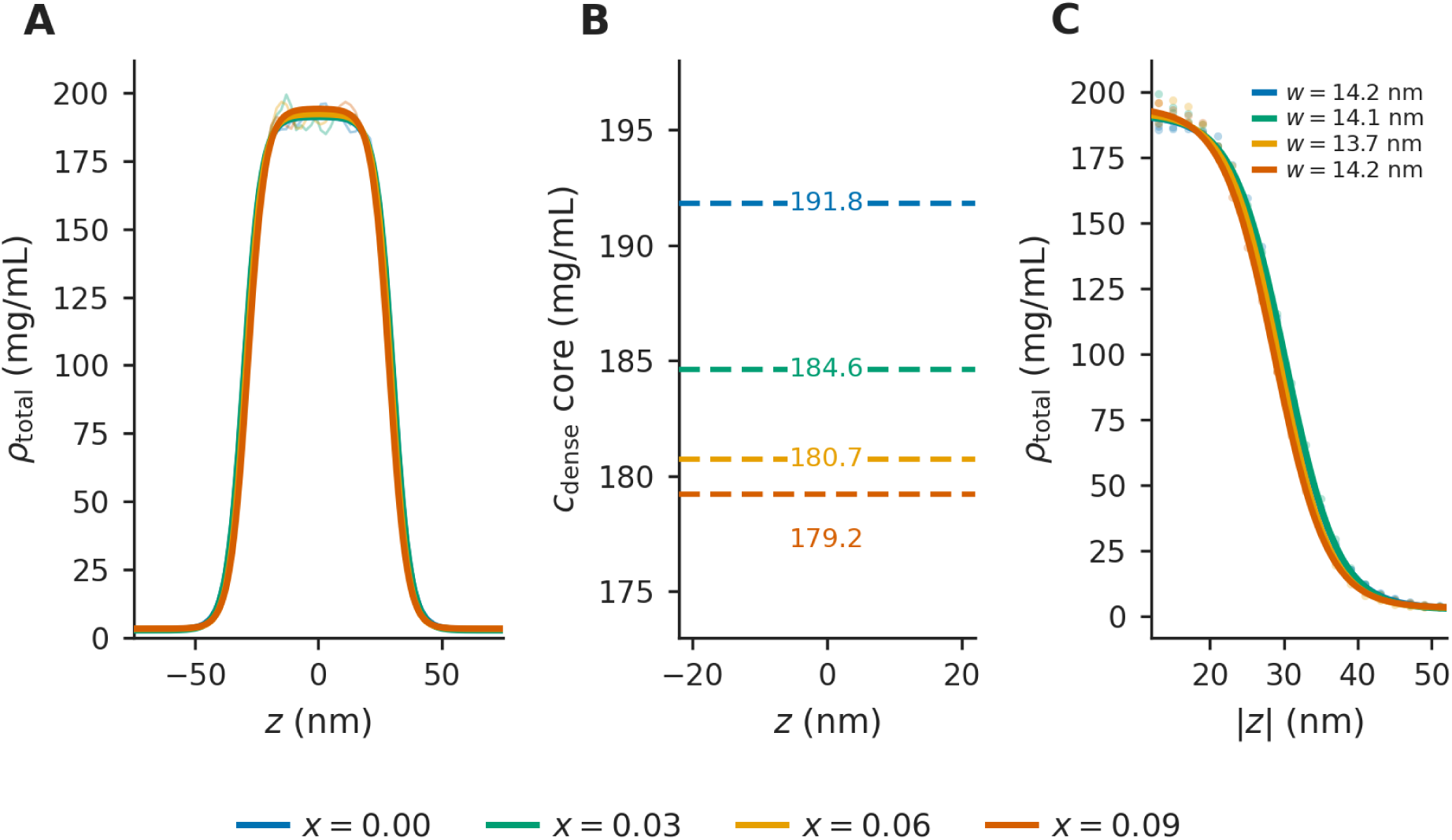
Slab density profiles and interface fits. Per-dose averages over the completed replicates at each HP6/Umbrea fraction *x*, colored by dose. (A) Total slab density *ρ*_total_(*z*) (all beads) with the max-gradient tanh interface fits. The total dense-phase profile remains essentially unchanged across dose, added HP6/Umbrea substitutes into freed volume. (B) The HP1a-probe dense-core density *c*_dense_ (the cohesive host network, |*z*| ≤ 10 nm), which falls monotonically with HP6/Umbrea (191.8 → 179.2 mg/mL, ∼7%, the pooled per-dose means of Fig 3A and S2 Table). HP6/Umbrea dilutes HP1a–HP1a contacts locally without dissolving the phase. (C) The interface folded to |*z*| with the tanh fits whose width *w* (≈13–14 nm, statistically flat across dose) sets *γ*.

**Fig S2.**
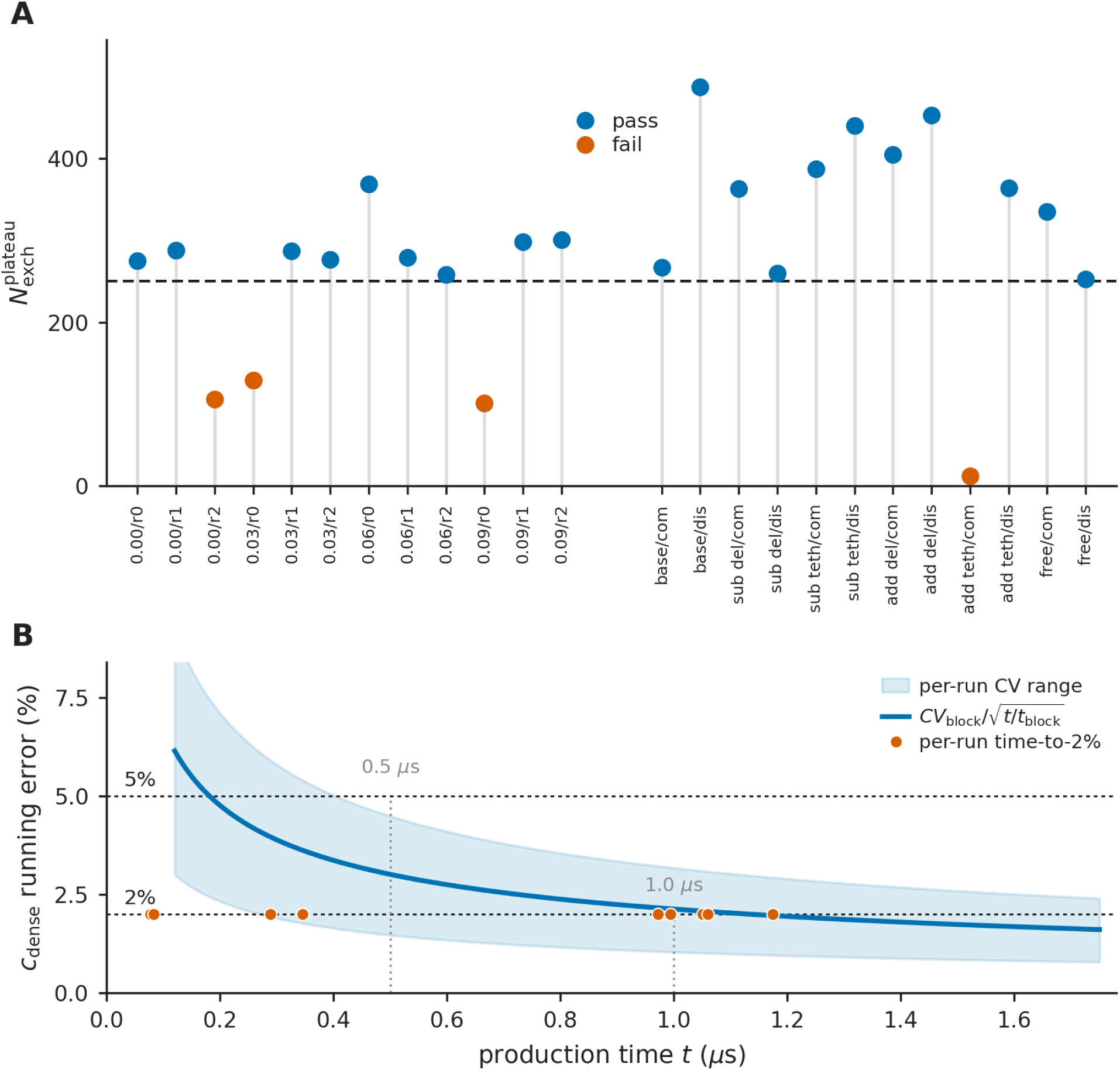
Convergence diagnostics. (A) Plateau exchange count per run for *N*_exch_ ≥ 250 gate (blue pass, vermillion fail). (B) Running convergence of *c*_dense_, per-run coefficient-of-variation band and the block-bootstrap 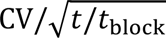 envelope reach ∼2% by ∼1 μs, supporting the two-level error model.

