## Supplementary figures and images for "HP6/Umbrea, a rapidly evolving *Drosophila* HP1-family paralog, is a candidate HP1a-recruited plasticizer of heterochromatin"

### Figure S1

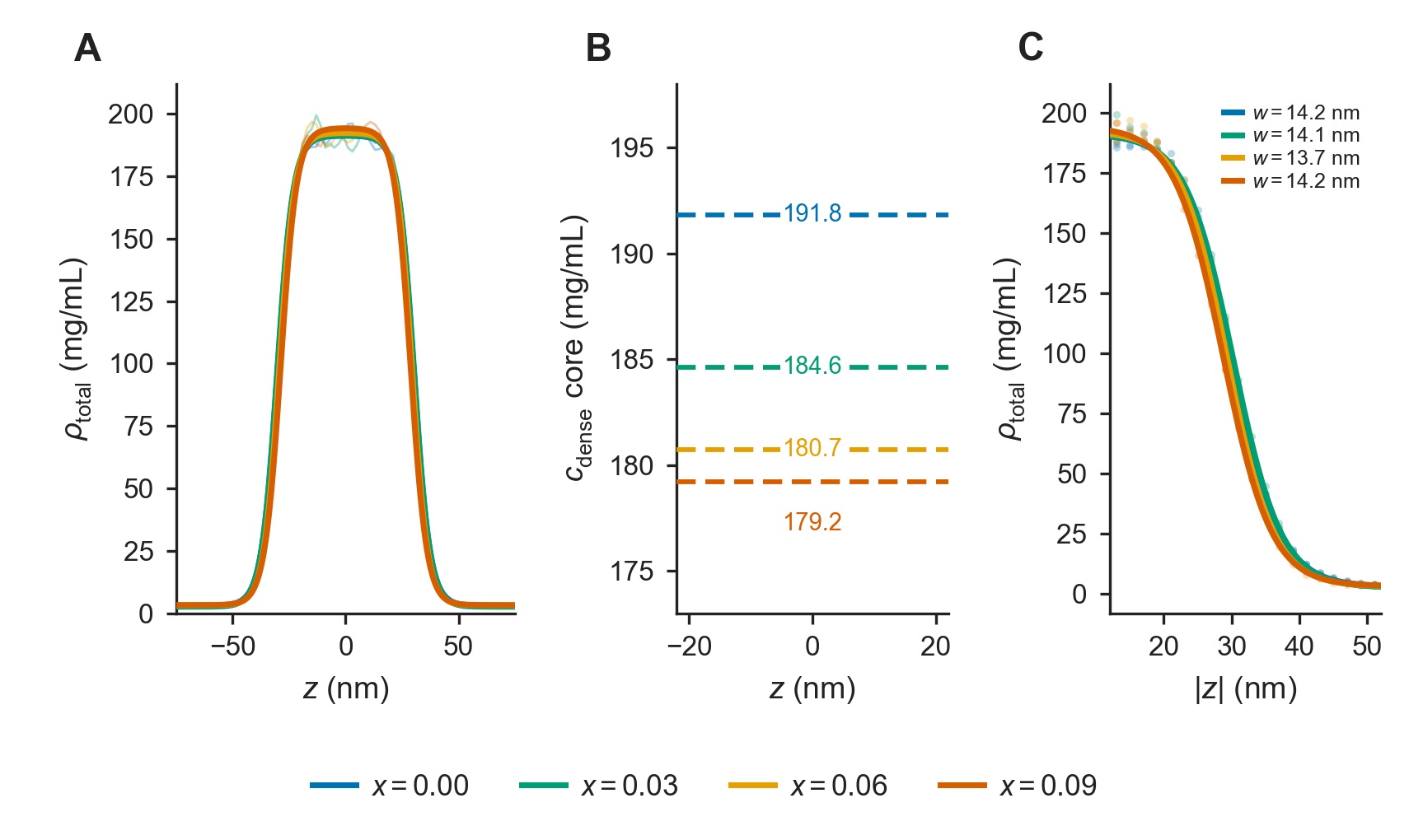

### Figure S2

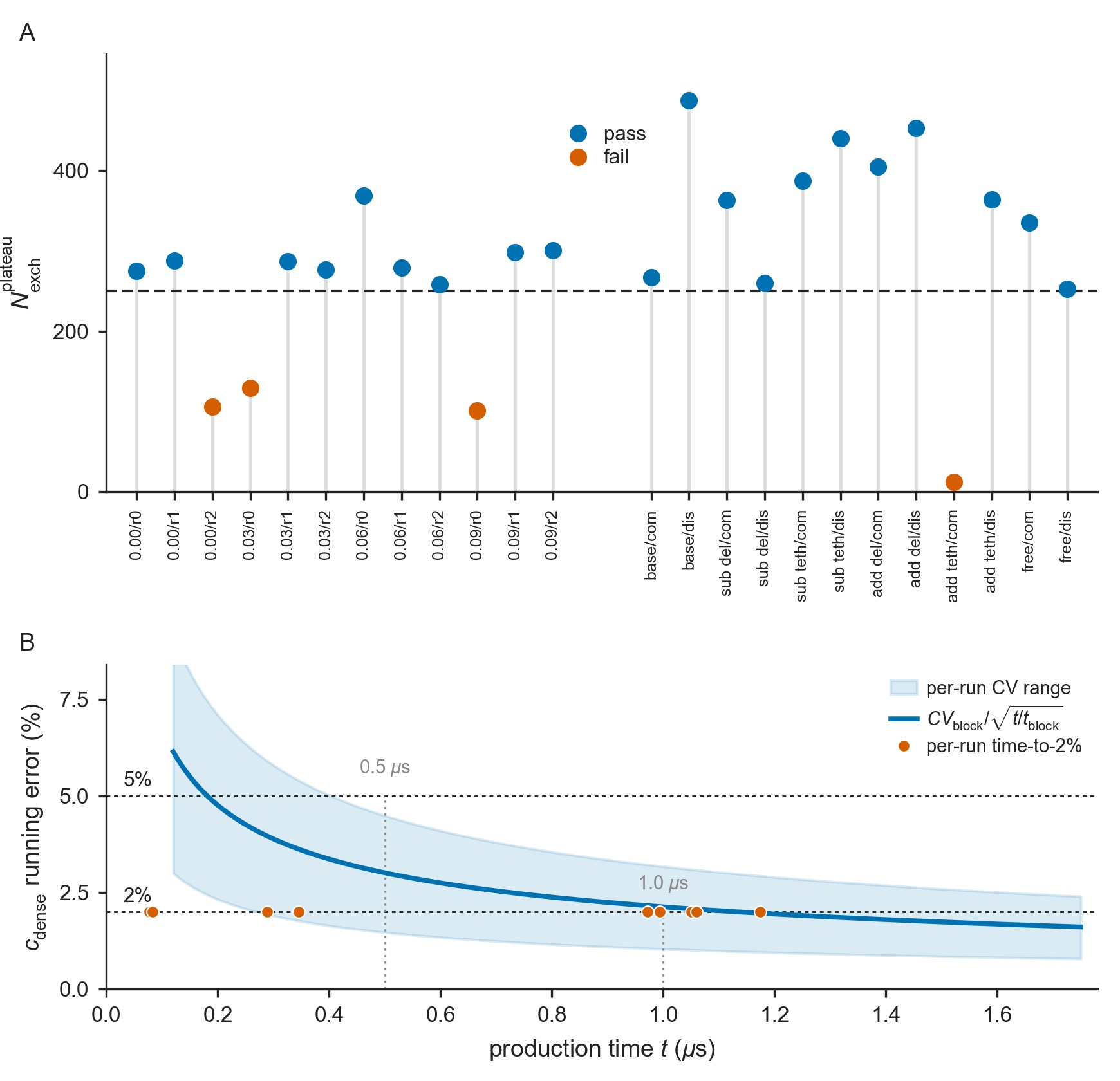
